# PELP: accounting for missing data in neural time series by Periodic Estimation of Lost Packets

**DOI:** 10.1101/2022.02.04.479162

**Authors:** Evan M. Dastin-van Rijn, Nicole R. Provenza, Wayne K. Goodman, Matthew T. Harrison, David. A. Borton

## Abstract

**Objective:** Implanted electrical stimulators with sensing capabilities have enabled the development of closed-loop neuromodulation therapies capable of responding to patient needs in real-time. Through a combination of rechargeable technologies and wireless data transmission, it is now possible for researchers to acquire extensive neural recordings from human participants in naturalistic settings using these bidirectional devices. However, data losses during wireless transmission hamper processing and the identification of neural signals of interest, driving the need for methodologies to properly estimate the impact of data loss.

**Approach:** To accurately reconstruct the timing of data containing losses, we have developed a method called Periodic Estimation of Lost Packets (PELP) to precisely determine the number of samples lost from implanted recordings during active stimulation. PELP leverages a data-driven procedure for determining the period of stimulation and the knowledge that stimulation continues identically during periods where data are missing to accurately account for the number of samples lost.

**Main results:** Using simulated stimulation added to collected human EEG data, we show that PELP is robust to a range of stimulation waveforms and noise characteristics. Lastly, we successfully applied PELP to local field potential (LFP) recordings from an implanted, bidirectional device using data recorded in the clinic and the patient’s own home.

**Significance:** By effectively accounting for the timing of missing data, PELP enables the analysis of complex, naturalistic neural time series data from bidirectional implanted devices aiding in the development of novel therapeutic approaches. NCT04806516 (ClinicalTrials.gov).

## 1. Introduction

Targeted electrical stimulation of the brain and spinal cord has proven to be a highly effective therapy for movement disorders, mental illnesses, and pain [1]. However, most neurological disorders do not have static symptoms while standard neuromodulation therapies rely on parameter updates during clinical visits that can be weeks to months apart [2–4]. This discrepancy leads to reductions in the efficacy of the therapy over time and an elevated risk of side effects from the stimulation [5]. One viable solution is adaptive, or closed-loop neuromodulation in which stimulation parameters are adjusted according to a known neural biomarker of disease state, thus tailoring therapies to a patients’ needs in real-time [2,5-7].

In order to aid in the development of closed-loop therapies, many implanted device manufacturers have designed ‘bidirectional’ implants capable of concurrently stimulating and sensing from the nervous system [8–11]. Early bidirectional devices stored data locally on the implant or required restrictive interfaces for wireless transmission of data limiting recordings to short time periods and unnatural environments. Furthermore, extensive sensing ran the risk of premature battery failure shortening device lifetime [12]. Recent investigational devices, for example the Medtronic Summit RC+S [8], BrainGate iBCI [13], and Neuralink [14] solve both of these limitations via rechargeable cell batteries and chronic, real-time streaming of neural data to external devices meters away. These advances allow access to long timescale neural recordings in natural environments, enabling the identification and development of personalized biomarkers and therapies [6,7,15,16].

During wireless transmission, neural data samples are often grouped into formatted units called ‘packets’ [17]. Packets contain a series of subsequent samples of a particular length as well as timing information and other relevant metadata. When transmitted, it is possible for packets to fail to reach the receiver leading to missing samples. These missing samples must be properly accounted for when the time series data is reconstructed. The timing information contained in each packet aids in this process but is frequently inexact due to hardware, network, and software delays [18], resulting in uncertainty in the number and location of the samples missing from a recording. This process, known as packet loss, is illustrated in Fig. 1a.

**Fig. 1.**
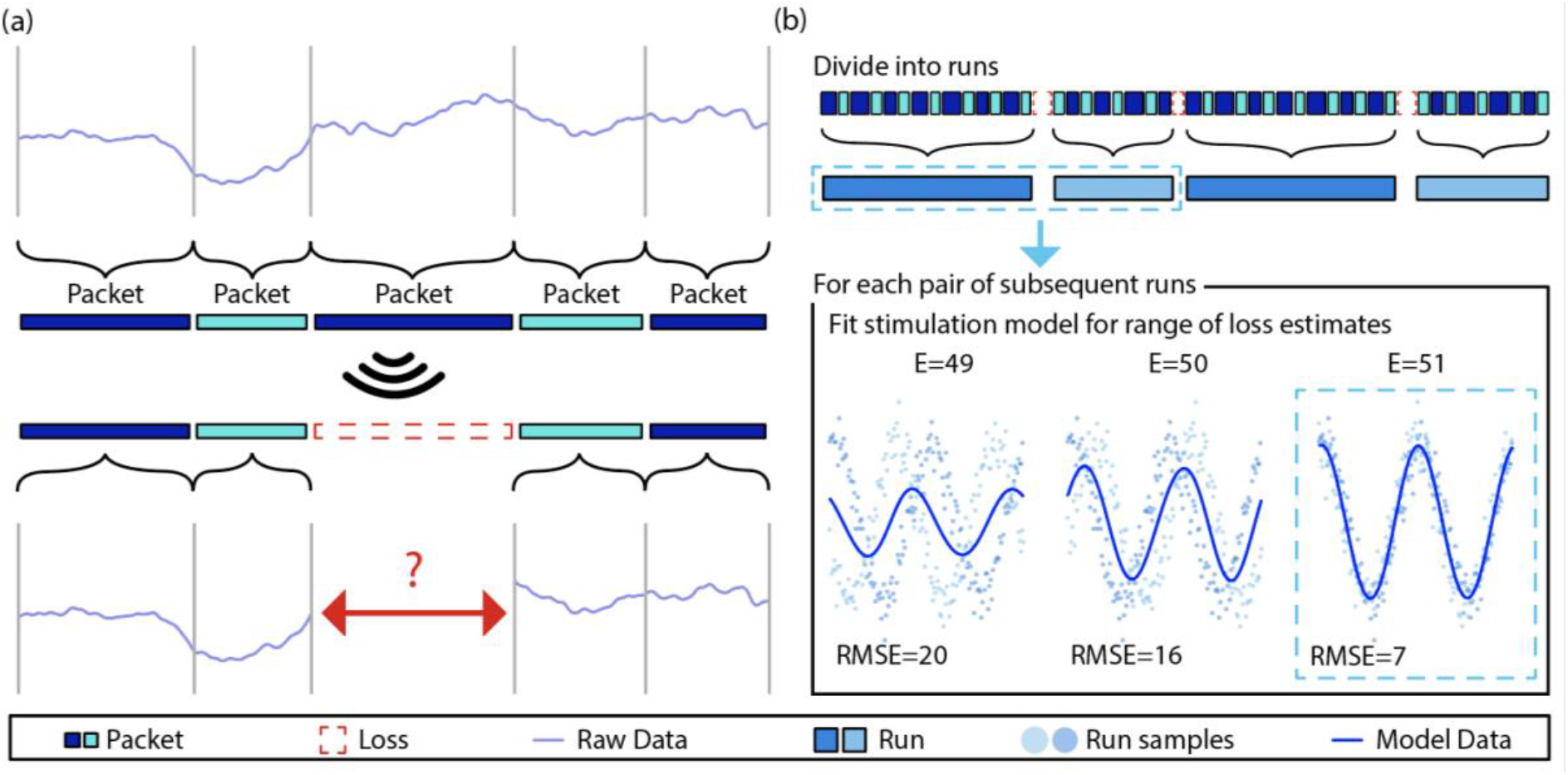
Illustration of packet loss and PELP. **(a)** Subsequent samples from a neural data time series acquired on an implanted device are grouped into packets. Packets are then wirelessly transmitted to a receiver. During the transmission process it is possible for some packets to be lost. As a result, the relative timing of the samples contained in received packets is uncertain. **(b)** PELP begins by grouping contiguous packets (blue, first row) into continuous runs (blue, second row) where each run is separated from adjacent runs by losses (dashed-red). Loss sizes are estimated but uncertain. The stimulation period is analytically determined using all the data. For each pair of subsequent runs, the root mean squared error (RMSE) between a stimulation model and the samples in the two runs is computed for a range of loss sizes centered around the estimate (indicated by E= for each size). A new stimulation model is fit for each loss estimate. The loss size that minimizes the RMSE is selected as the true loss size.

to a packet loss for recordings where stimulation is present. Before PELP can be applied to a recording, the locations of packet losses and their estimated sizes must first be determined. As an exemplar device, we focus our methodological development on the wireless data transmission from the Summit RC+S, however, the methodology is generally applicable. For recordings using the Summit RC+S, each packet has three integer timing variables of note for this purpose: ‘dataTypeSequence’ indicating the packet number that rolls over every 256 packets, ‘systemTick’ time of the last sample in a packet with 0.1 ms resolution that rolls over every 6.5536 s, and ‘timestamp’ with 1 s resolution and no rollover [22]. A packet loss has occurred when the dataTypeSequence between subsequent packets skips an index or the timestamps are inconsistent with the systemTick data. For sampling rate *f_s_* and *m* rollovers, the number of samples lost *N* can be estimated according to the following equation:

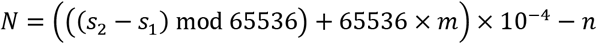

where *s_1_* and *s_2_* are the system ticks of the packets preceding and following the loss respectively and *n* is the number of samples in the packet after the loss. For loss segments greater than 6 seconds, the resolution of a systemTick is no longer acceptably accurate due to timing drift between the systemTick and timestamp. In these cases, the timing will need to be reset using a coarser metric such as the Unix timestamp (± 50 ms vs. ± 3 ms) corresponding to the time when the packet was received or generated. These estimates are not sufficiently accurate to ensure the exact reconstruction of the timing between received packets down to sample resolution. PELP leverages the presence of regular stimulation in both the received and missing data to ensure exact estimates of data losses.

Before applying PELP, we divide the time series into a set of *R* consecutive ‘runs’, where each run is composed of contiguous packets, and consecutive runs are separated by packet losses. The recording in run *r* is a sequence of *n_r_* (time,value) pairs ((*t_k,r_, y_k,r_*): *k* = 1,…, *n_r_*)). where *t_k,r_* is the time relative to the start of run *r* and *y_k,r_* is the recorded LFP amplitude at that time. Let *δ* be the period of stimulation, which we assume is constant across the entire recording. For regular stimulation, the stimulation artifact can be modeled as the *δ*-periodic function

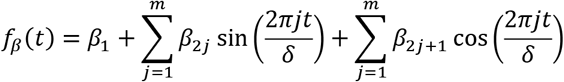

[23] for appropriate choice of the number of harmonics (*m*) and the parameter vector *β* = (*β_j_: j* = 1,…,2*m* + 1). Within a single run, *β* can be estimated via least-squares using the harmonic regression model *y_k,r_* = *f_β_*(*t_k,r_*) + *ϵ_k,r_* with homoscedastic noise *ϵ_k,r_*. Across multiple runs, however, the regression model will only be a good fit for appropriate choice of the packet loss sizes, a fact that we can leverage to estimate these loss sizes from data. Let Δ*_r_* denote the duration of the packet loss between runs *r* and *r* + 1. We estimate Δ*_r_* by choosing the one that gives the best least-squares fit to the harmonic regression model using the combined data from runs *r* and *r* + 1, i.e.,

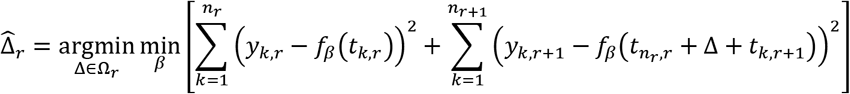

where Ω*_r_* is a (small) finite set of candidate loss sizes, in our case, the set of loss sizes corresponding to a positive integer number of samples, centered around the initial loss size estimate, and spanning the uncertainty in the estimate. The minimization over *β* can be done exactly for each Δ ∈ Ω*_r_* using least-squares, and then the optimal Δ can be selected. It is important that Ω*_r_* is based on an accurate initial estimate without too much uncertainty because candidate loss sizes that differ by an integer multiple of the period *δ* cannot be distinguished. The method is illustrated in Fig. 1b.

PELP requires knowledge of the stimulation period (*δ*) and an appropriate choice of the number of harmonics (*m*) used by the harmonic regression model. Although *δ* is known in principle, slight inaccuracies in device system clocks make it important in practice to use data-driven methods to estimate *δ* [23]. Before using PELP, we estimate *δ* from combined data across all runs using the multiple-channel period estimation method described in detail in [23], where we treat each run as a separate channel, where we use at most the first 10^4^ samples from each run, and where we use only the first two stages of the stagewise search for *δ* before the final optimization. We similarly choose *m* in a preprocessing step using Akaike Information Criterion (AIC) [24] for the harmonic regression model (with Gaussian errors) applied to the single longest run.

### 2.2. Participant

The research was conducted in accordance with the principles embodied in the Declaration of Helsinki and in accordance with local statutory requirements. The participant gave informed consent and data presented were collected in accordance with recommendations of the federal human subjects regulations and under protocol H-40255 approved by the Baylor College of Medicine Institutional Review Board. EEG data were recorded both with and without stimulation when the participant visited the clinic for DBS programming. LFP data were recorded both in the clinic and when the patient was at home. Electrodes on both the right and left sides were implanted in the VC/VS according to standard stereotactic procedures using computed tomography for target determination. The location of electrode placement was made entirely on clinical grounds. Bilateral 150.6 Hz stimulation with a pulse width of 90 μs and amplitude of 4 mA for the left side and 4.5 mA for the right side was used for all recordings where stimulation was turned on.

### 2.3. EEG and LFP Recording Procedures

Continuous electroencephalography (EEG) was recorded using a 64-channel ActiCap BrainVision system (Brain Vision, Morrisville, NC, USA). A common mode sense electrode was located at FCz. The EEG was band-pass filtered online between 0.1 and 1000 Hz and digitized at 5 kHz. The EEG was downsampled offline to 1000 Hz with an anti-aliasing filter prior to analysis. The continuous LFP was recorded using the Summit RC+S (Medtronic, Minneapolis, MN, USA) via wireless data streaming from implanted electrodes to the device running the task. Each DBS probe (Model 3387, one per hemisphere) contains four electrode contacts two of which were used per side to conduct bipolar recordings. LFP recordings were sampled at 1 kHz in the clinic and 250 Hz at home in order to minimize data losses. Signal processing and analysis were performed in Matlab (Mathworks, Natick, MA, USA) using in-house code.

### 2.4. Stimulation Simulation Procedure

In order to simulate stimulation in our recordings, we modeled DBS artifacts as a sum of sinusoidal harmonics of the stimulation frequency [25]. The effect of stimulation was simulated by adding the artifact component regressed from recordings on a different day where stimulation was turned on to data without stimulation. A high-pass filter at 1 Hz with a gaussian window was first applied to achieve approximately 40 dB attenuation in the stopband before the period of stimulation (*δ*) was identified using the period estimation component of PARRM [23]. A sum of sinusoids *f_β_*(*t*)with *m* harmonics of the period and coefficients *β* was then fit to the data using linear regression. The stimulation amplitude for each cycle was sampled from a normal distribution with mean A_1_ and standard deviation *v*. The mean stimulation amplitude was set relative to the root mean squared amplitude of the stimulation off data (A) according to a ratio (R) and the original root mean squared amplitude of the fit (A_o_). In order to model potential inaccuracies in period estimation, the period of the stimulation model was slightly offset by a drift factor (d) measured in percent drift per 1000 cycles from the period used during PELP. The effects of these three parameters are illustrated in Fig. 2.

**Fig. 2.**
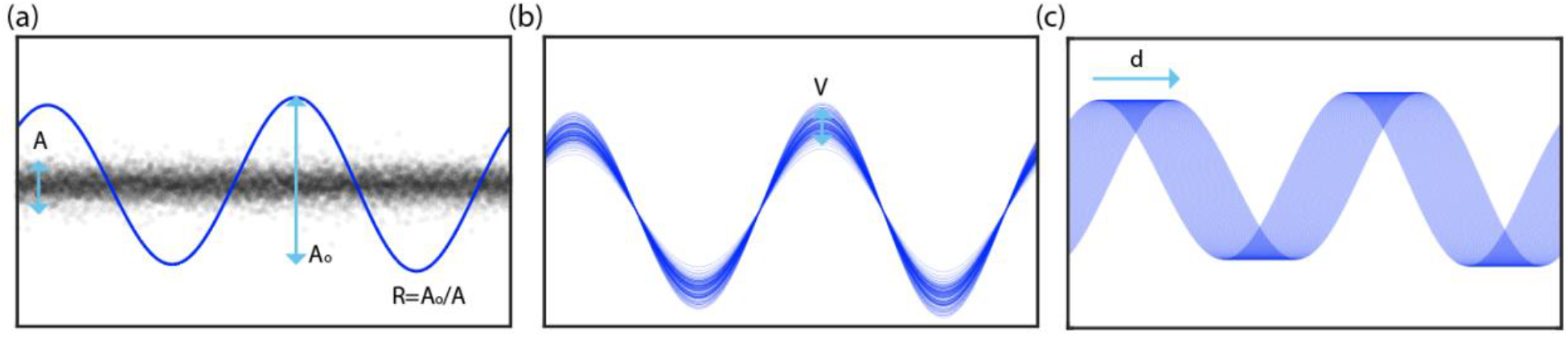
Illustration of simulation components. **(a)** The root mean squared amplitude (Ao) of the stimulation model is set relative to that of the neural signal of interest (A) according to a target ratio R. **(b)** The amplitude of each stimulation pulse is varied on a cycle-wise basis where the amplitude of each pulse is sampled from a normal distribution with mean Ao and standard deviation V. **(c)** Inaccuracies in period estimation, drifting sampling rate, and frequency variability are modeled by adding a drift factor d to the stimulation period in the model.

### 2.5. Computational Experiments

We conducted three sets of experiments to simulate the accuracy of loss estimation while varying different parameters in the stimulation model. For each, Monte Carlo analyses were used to simulate a large number of experiments by randomly sampling subsets of 50 sample “packets” to remove from the recording. These simulations were applied while varying one of amplitude ratio, amplitude variability, or drift as a function of the loss uncertainty. For each simulation, the uncertainty ranged from 0-50 samples in 1 sample increments while the dependent parameters ranged from 0-4 in increments of 0.1 for the amplitude ratio, 0-10% in increments of 1% for the amplitude variability, and 0-0.6% in increments of 0.015% per 1000 cycles for the drift. PELP was applied to each simulation to determine the proportion of losses it was able to estimate correctly depending on the stimulation parameters. This approach is similar to that of Boudewyn et al. [26] and is informative because it uses a combination of real EEG data similar to LFP (so that the noise properties are realistic) and artificially induced losses (so the actual truth is known) across a range of modeled stimulation waveforms.

## 3. Results

### 3.1. Stimulation Model Fit

We first sought to estimate the frequency of stimulation to construct a model for time series data alignment. Fig. 3a shows the stimulation model compared to the raw EEG samples overlapped by computing the modulo of each timepoint with the model’s period of stimulation. The period of stimulation was found to be 6.64000 samples. Four sinusoidal harmonics were used for the fit based on model selection via AIC. Raw samples are well consolidated about the artifact waveform with a residual standard deviation of 5.90 μV similar to the standard deviation of the stimulation off data (4.55 μV).

**Fig. 3.**
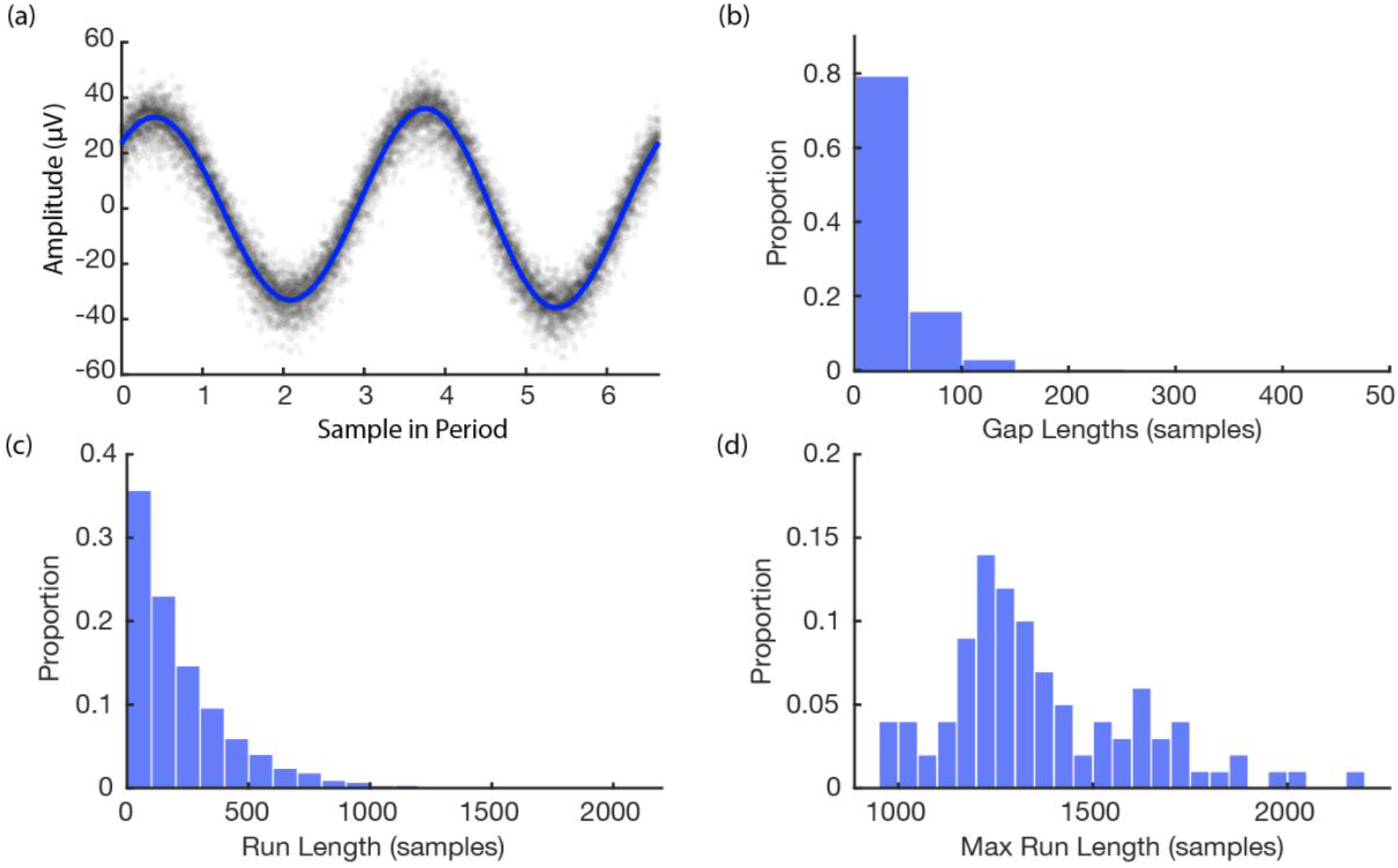
Features of simulation. **(a)** Stimulation model fit to EEG data. Raw data are shown in gray and the model is shown in blue. **(b)** Histogram of missing data gap lengths for all experiments. **(c)** Histogram of continuous run lengths for all experiments. **(d)** Histogram of longest continuous run in each experiment

### 3.2. Loss Simulations

After building the model of stimulation times, we then ran a set of simulations to provide bounds on the expected recovery and overall loss after using the PELP method for data alignment. The histograms in Fig. 3 illustrate features of the Monte Carlo simulation of 100 loss experiments. The average length of a missing data gap was 63 samples with a median of 50 samples (one loss). Runs of continuous samples between losses ranged from 50-2200 samples with an average length of 251 samples. The max run length in each simulation ranged from 950-2200 samples with an average length of 1342 samples corresponding to roughly 202 cycles of the 150.6 Hz simulated stimulation frequency.

### 3.3. Loss Estimation Experiments

We then explored the impact of model parameters on the loss of sensing data using PELP. Fig. 4 shows the Monte Carlo simulated loss experiments measuring the accuracy of PELP estimates as a function of the stimulation amplitude ratio, amplitude variability, and estimate uncertainty. Both heat maps show discrete transitions in accuracy at uncertainty multiples of three samples. This occurs since for bilateral stimulation with a period of roughly 6.64 samples, estimated differences of multiples of three samples will be more closely overlapping. For constant uncertainty, accuracy increased smoothly for increasing amplitude ratio and decreasing amplitude variability. Changes in drift, in the range tested, had no effect on accuracy. Keeping amplitude ratio or amplitude variability constant while varying uncertainty had little effect on accuracy with exception of the effects at multiples of three samples. Changes in uncertainty for constant drift had no effect on accuracy. Accuracy was near 100% for amplitude ratios above 0.2 for uncertainties less than 3 samples, amplitude ratios above 0.5 for uncertainties less than 9 samples, and amplitude ratios above 3 for uncertainties greater than 9 samples. Accuracy was near 100% for amplitude variabilities below 2% for uncertainties less than 9 samples. For uncertainties larger than 9, amplitude variability had to be near zero to maintain 100% estimate accuracy. Across all values for the drift experiment, accuracy was near 100%.

**Fig. 4.**
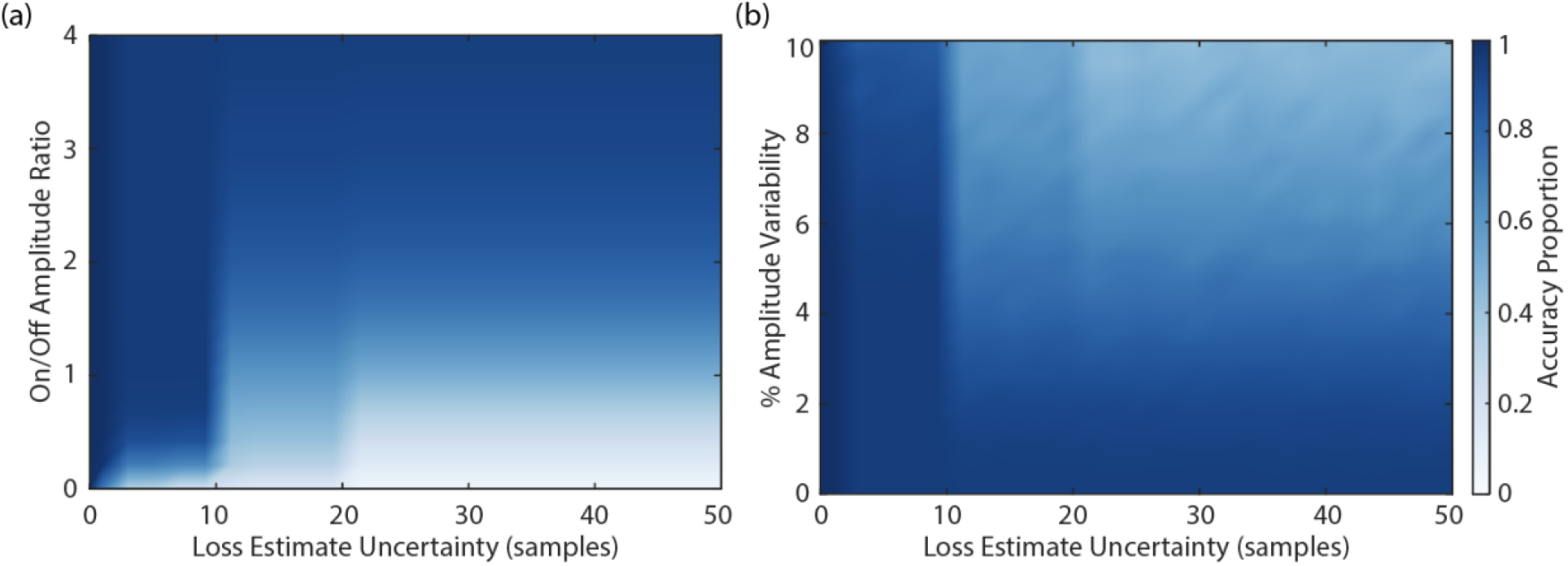
Accuracy of loss estimation as a function of amplitude ratio, amplitude variability, and uncertainty. The accuracy of loss estimation was computed for 100 simulated trials with 20% of the packets removed. More accurate parameter combinations are indicated by darker values in the colormap. Amplitude ratios **(a)** ranged from 0-4, amplitude variability **(b)** ranged from 0-10%, and uncertainty ranged from 0-50 samples.

### 3.4. PELP with Summit RC+S Recordings

We then applied the PELP methodology of offline data realignment to brain recordings collected from participants of an ongoing clinical study to demonstrate real world performance. Figure 5 shows LFP data from a behavioral task containing packet losses recorded using the Summit RC+S in the clinic and at-home after estimation of losses using PELP. Data recorded in the clinic sampled at 1000 Hz contained 15 losses with a median size of 200 samples (Fig. 5a-b). Data recorded at home sampled at 250 Hz contained 121 losses with a median size of 17 samples (Fig. 5c-d). When overlapped on the timescale of the period of stimulation, all samples from both conditions were well consolidated about the stimulation waveform with no observable evidence of significant period drift indicating that losses were accurately accounted for.

**Fig. 5.**
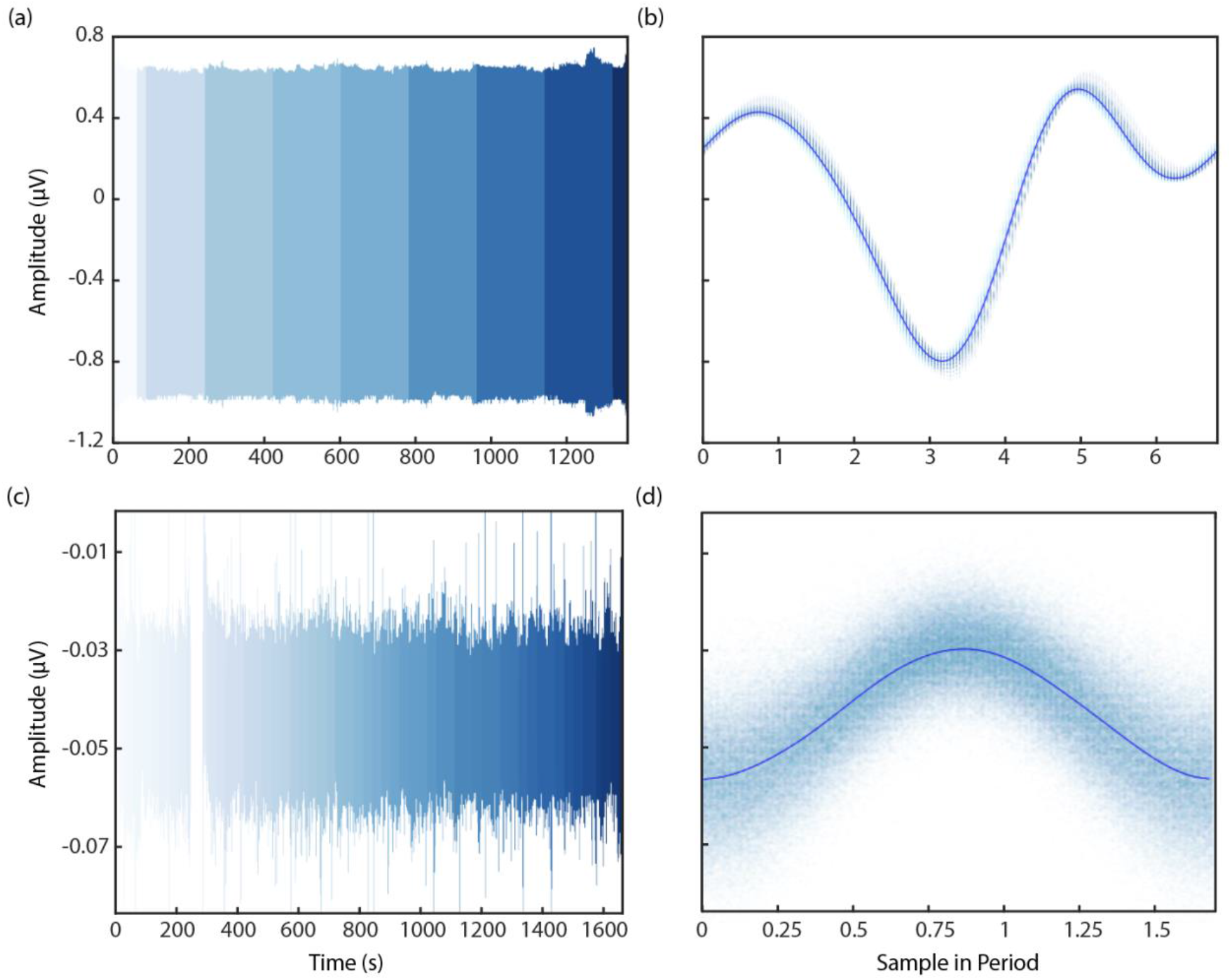
Application of PELP to data from the Summit RC+S. PELP was applied to RC+S data containing losses sampled at 1000 Hz **(a, b)** and 250 Hz **(c, d)**. Each continuous run is indicated by a distinct shade in the colormap **(a, c)**. For both conditions, samples from all runs were overlapped on the timescale of the period of stimulation **(b, d)**. Samples were well consolidated about the stimulation model for both conditions indicating accurate estimation of loss sizes.

## 4. Discussion

Streaming of intracranial electrophysiology data in the clinic and at home in ecologically valid environments is essential for biomarker discovery in a variety of neurological disorders. Bidirectional implanted devices have enabled the acquisition of such datasets, however, data losses during wireless streaming hinder accurate analyses of neural signals. To address these issues, we have developed PELP to exactly estimate and account for data losses from implanted recordings where stimulation is on. We show using simulations of data losses that PELP can accurately estimate missing samples over a variety of stimulation conditions. Lastly, we successfully applied PELP to reconstruct the timing of data recorded using the Summit RC+S in the clinic and in the participant’s home.

PELP is widely applicable to other stimulating devices capable of wireless data streaming. Our stimulation model accurately accounts for the range of amplitude and variability parameters that could be expected for other implanted devices. In recordings where sensing and stimulation occur on nearby contacts, stimulation amplitude can exceed the underlying neural signal by a factor of 10 [27]. For recordings where sensing and stimulation occur far apart or the stimulation harmonics fall within the transition band of an online low-pass filter, the amplitude ratio will be closer to 1. Our recordings using the Summit RC+S had amplitude ratios of 27 and 1.2 and estimate accuracy was consistent with predictions from the simulation. Pulse to pulse amplitude variability for the Summit RC+S is well within the range of values where PELP was most accurate. Fluctuations in battery or the surrounding medium could influence amplitude on longer timescales. While only the run nearest to the loss was used for estimation, drift within the run itself would not be well accounted for. Exceptionally long runs or runs where drift was identified could be divided to improve estimation accuracy. Similar considerations would also be effective if stimulation frequency drift or errors in period estimation occur.

Since PELP requires stimulation artifacts to be present in order to model the signal during data losses, the method is not applicable for recordings where stimulation is off or significantly attenuated by online filters. In such circumstances, less accurate methods utilizing packet timing metadata must be used for loss estimation. In theory, stimulation could be applied below therapeutic amplitudes and still be used for reconstruction using PELP although such modifications would only be reasonable if the inevitable stimulation artifacts did not obscure neural signals of interest.

Additionally, in the case of discontinuous bursts of stimulation PELP could be adapted to assist in artifact removal. Bursts of stimulation are useful both clinically for the treatment of neurological disorders like epilepsy [28] and in research for investigating many parameter combinations [29], however, their brevity inhibits the use of template subtraction-based methods for artifact removal. PELP could be used to calculate differences in phase between separate bursts of stimulation with the same parameters thereby enabling the creation of a combined dataset with aligned stimulation pulses. Such an approach could be useful for increasing the amount of data available to artifact removal methods relying on template subtraction.

## 5. Conclusion

By leveraging the periodicity of stimulation artifacts, PELP produces highly accurate reconstructions of timing information from wirelessly transmitted neural data. PELP effectively accounts for the timing of missing data enabling the analysis of complex, naturalistic neural time series data from existing and next-generation bidirectional implanted devices aiding in the development of novel therapeutic approaches.

